# Single-dose neonatal gene therapy preserves motor coordination and extends survival of *Polg*-deficient mice

**DOI:** 10.64898/2026.11.17.762423

**Authors:** Sofia Zanin, Stéphanie Moriceau, Elissa Afram, Maria Teresa Pérez-Cartier, Clovis Gaborit, Marcelo Simon Sola, Stephen C. Kneeland, Stephen A. Murray, Arnold Munnich, Benedetta Ruzzenente, Metodi D. Metodiev

**Affiliations:** Imagine Institute, Université Paris Cité, INSERM UMR1163, Paris, France; Neurobehavior and Metabolism Platform, Institut Imagine, Structure Fédérative de Recherche Necker, INSERM US24/CNRS UAR, Paris, France; AAV platform, Imagine Institute, Paris, France; The Jackson Laboratory, Bar Harbor, Maine, USA

**Keywords:** POLG, gene therapy, mitochondrial disease, adeno-associated virus

## Abstract

*POLG* mutations are the most common single nuclear gene cause of mitochondrial disease, resulting in a broad spectrum of clinical presentations with varying ages of onset and disease severity. POLG disease is incurable, with few symptomatic treatments available that cannot prevent its progression. Gene replacement therapy has the potential to halt or prevent POLG disease. Here, we inactivated the *Polg* gene in mouse parvalbumin-expressing neurons and fast-twitch skeletal muscle fibers. Knockout mice developed motor coordination deficits, respiratory chain deficiency in Purkinje cells and skeletal muscle, and died prematurely. In parallel, we developed nine AAV-based gene therapy vectors and confirmed their functionality *in vivo*. Finally, a single high-dose administration of two lead vector candidates into *Polg* KO neonates preserved respiratory chain activity in Purkinje cells and skeletal muscle, delayed disease onset, and extended lifespan. Our work provides preclinical proof of concept for gene replacement therapy for POLG disease, laying the foundation for further development in the field.

## Introduction

*POLG* is a nuclear gene encoding POLγA, the 140 kDa catalytic subunit of the human POLγ enzyme. The latter, together with the TWINKLE helicase and the single-strand binding protein SSB, forms the so-called minimal replisome that, in concert with other factors (POLRMT, TFB2M, etc.), replicates mitochondrial DNA (mtDNA) ^1–3^. POLγ is a heterotrimer composed of one POLγA subunit and two copies of the smaller auxiliary subunit, POLγB (encoded by *POLG2*). POLγA carries the DNA polymerization, 3′–5′ exonuclease proofreading and 5’-deoxyribose-phosphate lyase activities of the complex, whereas POLγB stabilizes the catalytic subunit and increases the DNA binding and processivity of the POLγ enzyme ^4,5^. Pathogenic mutations in either *POLG* or *POLG2* can impair mtDNA replication, leading to mtDNA depletion or multiple deletions, respiratory chain (RC) deficiency, and primary mitochondrial disease (PMD) ^6,7^.

The *POLG* gene is the most frequently implicated nuclear gene in PMD. Population genetic analyses estimate the lifetime risk of autosomal recessive POLG disease at ∼1 in 27000, whereas a nationwide Norwegian study reported a birth prevalence of 1 in 48000 ^8,9^. To date, more than 300 disease-causing variants have been mapped to the *POLG* locus, including missense, nonsense, splice-altering and indel variants with recessive or autosomal dominant modes of inheritance (https://portal.niehs.nih.gov/polg/). It is estimated that as many as ∼2% of the population may carry *POLG* disease variants ^6^. Variant distribution along the *POLG* sequence is nearly uniform, with limited genotype-phenotype correlation. Generally, autosomal dominant *POLG* variants are associated with relatively milder late-onset pathology (e.g., progressive external ophthalmoplegia (PEO), myopathy, ataxia), whereas autosomal recessive variants often cause earlier-onset and more severe, often fatal, disease (e.g., Alpers-Huttenlocher syndrome (AHS), myocerebrohepatopathy (MCHS)) ^10^. To establish a basis for phenotype-genotype correlations in POLG disease, Farnum and colleagues assigned known disease-causing variants to functional clusters based on their structural locations and predicted biochemical effects ^11^. This approach revealed that compound heterozygous mutations assigned to different POLγA clusters present with an earlier onset and more severe disease than homozygous recessive mutations or compound heterozygous mutations within the same cluster. Clinically, POLG disease is often multisystemic and spans a spectrum of clinical presentations with variable onset and severity ^10,12^. In neonates with POLG disease, the presence of liver involvement and/or epilepsy progressing to status epilepticus is associated with increased mortality ^13–15^.

POLG disease is incurable. Available treatments are supportive and aimed at symptom management. They may include mitochondrial cocktails containing CoQ10, riboflavin, thiamine, and antioxidants, or a ketogenic diet to control seizure frequency ^6^. However, they cannot stop disease progression. In contrast, gene replacement therapy, which directly targets the genetic cause of the disease, can be curative. In this approach, a functional copy of the *POLG* coding sequence is introduced using recombinant adeno-associated viruses (rAAVs) as delivery vehicles. However, until recently, research in the field was blocked by the large size of the *POLG* cDNA, which was considered too large to fit into a classical rAAV genome, and by the lack of suitable animal models for preclinical development. The *POLG* cDNA is 3.7 kb and, when combined with standard regulatory elements, could easily exceed the limited cargo capacity of classical rAAV vectors (4.9-5 kb) ^16^. While it is possible to package large genomes into AAV capsids, this is associated with a steep decrease in production titers and a risk of rAAV genome truncation ^16^. In other cases, large coding sequences have been successfully delivered using a split-vector approach, in which the insert is divided into two fragments, each carried by an rAAV plasmid ^17,18^. However, because this approach relies on co-transduction of the same cell with both rAAV vectors, it generally results in reduced transgene expression and partial genomes in individual cells ^19^. Recently, we have successfully used size-optimized vectors carrying minimal regulatory elements to deliver the even larger *LRPPRC* cDNA into different mouse tissues, partially overcoming the size-limitation roadblock ^20^.

Another roadblock to developing *POLG* gene replacement therapy was the lack of suitable animal models with a robust disease-relevant phenotype. Animal models continue to play an important role in preclinical development of *in vivo* gene replacement therapies because, unlike in vitro systems, they can recapitulate the biodistribution and transgene expression of viral vectors across different tissues. At the outset of this project, the available mouse models included the so-called mtDNA mutator mouse ^21^ and a knock-in model expressing the mouse equivalent of the common recessive A467T disease-causing variant ^5^. The mutator mouse expressed the proofreading-impaired POLγA^D257A^ variant, causing accumulation of mtDNA mutations and a premature aging phenotype. Neither the variant nor the phenotype has been reported in POLG disease. The A467T knock-in mice presented mild molecular and behavioral phenotypes but appeared otherwise healthy and had a normal lifespan.

Here, we demonstrate proof of concept for *POLG* gene replacement therapy. First, we developed a conditional knockout mouse model in which we inactivated the *Polg* gene in parvalbumin-expressing (PV+) cells, resulting in progressive motor coordination deficits, RC dysfunction in Purkinje cells and skeletal muscle, and a shortened lifespan. In parallel, we generated nine gene therapy vectors with different promoter-*POLG* cDNA combinations. *In vitro* and *in vivo* analyses demonstrated that all nine vectors transduced various mouse tissues, including the central nervous system (CNS), and drove immunodetectable expression of human POLγA protein. Next, we injected two vectors carrying the wild-type and codon-optimized *POLG* cDNA under the *EFS* promoter into the temporal vein of newborn KO mice. This single-dose injection delayed disease onset, preserved RC function in Purkinje cells and skeletal muscle, and extended the lifespan of treated KO mice, demonstrating the therapeutic potential of our gene replacement vectors. To our knowledge, this is the first proof-of-concept study for gene replacement therapy for POLG-linked disease, one of the most frequent mitochondrial diseases.

## Methods

### Mouse model, housing and experimentation

Conditionally targeted *Polg^Lox/Lox^* mice were generated at The Jackson Laboratory (JAX) (C57BL/6NJ-Polg^em1Murr/Mmjax^, strain no. 072088, available at the MMRRC) using CRISPR/Cas9 gene editing (guide RNA sequences 5′-GTGGGAGGCGAATAGTAAAG-3′ and 5′-CTGTCTTCCCTAAAGACCGC-3′). *LoxP* sites were inserted upstream and downstream of exon 3 of the *Polg* gene in C57BL/6NJ zygotes that were then transferred to pseudopregnant females, and *LoxP*-carrying progeny were identified by Sanger sequencing of the target region. To obtain germline transmission of the *Polg^Lox^ allele*, male progeny were bred to C57BL/6NJ females. To generate a tissue-specific *Polg* knockout in PV+ cells, *Polg^LoxP/LoxP^* mice were crossed to mice carrying *Cre* recombinase under the control of the parvalbumin promoter obtained from JAX (PV-Cre, B6.129P2-*Pvalb^tm^*^1^(cre)*^Arbr^*/J, strain no. 017320). Knockout mice had the genotype *Polg^Lox/Lox^, PV-Cre^+/-^* (henceforth KO mice), whilst controls were *Polg^Lox/Lox^* (henceforth L/L). Wild-type C57BL/6NCrl mice were purchased from Charles River.

To map *PV-Cre*-mediated recombination, *PV-Cre* mice were crossed to the reporter line *Rosa26^tdTomato^*, which carries a *Cre*-inducible tdTomato transgene ^22^, kindly provided by Dr. Alessandra Pierani (Imagine Institute, Paris).

Mouse housing and experimentation adhered to the European Union and French animal welfare legislation (Directive 2010/63/EU), and all mouse procedures were approved by the local institutional ethics committee (CEEA 34) and the French Ministry for Higher Education and Research (#45929-2023012016066698 v18). Mice were housed in rooms under controlled conditions (20–22°C, 45–64% humidity) on a 12 h light/dark cycle and fed a standard diet *ad libitum*.

Temporal vein injections were performed on neonates at P0–P1 ^23^. Prior to injection, mice were anesthetized by hypothermia on ice for ∼60 seconds. 1–2 × 10^14^ vector genomes/kg were injected in a total volume of up to 30 μl using a Hamilton glass syringe equipped with a 31G needle (10 mm, point type 4). At birth, sex and genotype cannot be determined immediately. Therefore, both L/L and KO neonates, regardless of sex, were injected with AAV vectors.

Body weight (BW) was recorded weekly. Relative body weight change with age was calculated as the ratio of the body weight at a given age to the body weight at the start of the experiment.

Adult mice were sacrificed either by cervical dislocation or by transcardiac perfusion with PBS under deep anesthesia induced by intraperitoneal injection of ketamine (180 mg/kg) and xylazine (10 mg/kg). Depending on downstream analyses, tissues were either snap-frozen in liquid nitrogen (for molecular biology experiments), frozen in isopentane (skeletal muscle for histology), or embedded fresh in OCT (brain for COX and SDH staining) and then frozen in liquid nitrogen. For immunofluorescence, brains were fixed overnight in 4% paraformaldehyde in PBS, cryoprotected by incubation in 20% sucrose overnight and embedded in OCT. For hematoxylin and eosin staining, brains were fixed overnight in 4% paraformaldehyde in PBS and embedded in paraffin.

### Construction of gene therapy plasmids and viral production

The human *POLG* cDNA (NM_002693.3) is 3.7 kb excluding the 5’- and 3’-UTRs. Wild-type *POLG* cDNA (*POLG^WT^*), codon-optimized (*POLG^CO^*) and CpG-depleted (*POLG^dCpG^*) variants fused to a C-terminal *FLAG* tag were synthesized and subcloned into a pUC57 vector by Genecust (Boynes, France). Codon optimization was carried out by Genecust, whereas CpG depletion was performed in-house by synonymous substitution of wild-type codons to disrupt CpG dinucleotides. Next, *POLG^WT^*, *POLG^CO^* and *POLG^dCpG^* were cloned into the *NotI*–*SpeI* sites of reduced-size rAAV plasmids carrying either a minimal cytomegalovirus (miniCMV) or a human elongation factor 1 alpha short form (*EFS*) promoter ^20^, yielding TGPOLG1-6. To generate plasmids TGPOLG7-9, the *EFS-intron* cassette from TGPOLG4-6 was replaced with a synthetic *CAG* (CMV enhancer fused to chicken beta-actin promoter)*-intron* cassette using the *AflIII/NheI*–*XbaI* restriction sites. The FLAG-free variants, TGPOLG4.1 and TGPOLG5.1, were generated by PCR amplification of the *POLG^WT^* and *POLG^CO^* cDNA with a downstream primer lacking the FLAG-tag sequence (*POLG^WT^*, upstream primer: 5′-ACCAATAGAAACTGGGCTTGTCG-3′, downstream primer: 5′-CTGACTAGTTCATGGGCCGGGCTGG-3′; *POLG^CO^*, upstream primer: 5′-CTAGAGCGGCCGCGCCACCATGAGC-3′, downstream primer: 5′-CTGACTAGTTCATGGGCCGGGCTGG-3′) and cloned into plasmid backbones carrying an *EFS* promoter via the *NotI*–*SpeI* sites. The TG7-GFP plasmid, containing GFP controlled by an *EFS* promoter, was described previously ^20^.

AAV vectors were produced by transient transfection in HEK293 cells (ATCC) and purified using CsCl-gradient centrifugation. Viral titers were determined by qPCR to measure viral DNA and by SDS-PAGE, followed by SYPRO Ruby staining and densitometric analysis.

### Rotarod analysis

Motor coordination and motor learning were evaluated using an accelerating rotarod (Rotarod/RS, Bioseb, France) ^24^. On day one, mice underwent 2 habituation trials of 120 s (acceleration from 4 to 20 rpm over 120 s) followed by 3 training trials of 5 min (acceleration from 4 to 40 rpm over 5 min). On day 2 and in the following weeks, mice underwent 3 trials with an acceleration from 4 to 40 rpm over 5 min. The dependent measure was the average time each mouse remained on the rotating rod before falling, defined as the latency to fall (LTF).

### Viral genome copy number and gene expression analysis

To measure viral genome copy numbers (VGCN), DNA was extracted from pulverized tissues using the DNeasy Blood and Tissue kit (Qiagen) following the manufacturer’s instructions. VGCN was determined using a QX200 droplet digital PCR system (Bio-Rad) with primers specific to *POLG^WT^* (upstream primer: 5′-TGGCAAACGGTAGAAGAACT-3′, downstream primer: 5′-AGTCAGAGCTAGGGGTTGA-3′), *POLG^CO^* (upstream primer: 5′-CAAAATGATTTCCTTCTGGAGAAAC-3′, downstream primer: 5′-GTCGTAGTCGGGGTGTC-3′) and *POLG^dCpG^* (upstream primer: 5′-AGTTCTTCTACCGTTTGCA-3′, downstream primer 5′-AATCTGGCTAAGCTGCCTAC-3′). Vector genome copies per diploid cell were calculated as twice the ratio of the vector concentration to the *Actb* concentration (upstream primer: 5′-CATCACTATTGGCAACGAGC-3′, downstream primer 5′-CAGGAGTTTTGTCACCTTTAGAT-3′), assuming two copies of the *Actb* reference locus per diploid cell.

To measure POLG transgene expression, RNA was extracted from pulverized tissues using TRIzol (Thermo Fisher Scientific) and the miRNeasy kit (Qiagen), with DNase treatment to remove contaminating DNA. cDNA was synthesized using the QuantiTect Reverse Transcription Kit (Qiagen). POLG mRNA was quantified by SYBR Green RT-qPCR using the primers described above. Normalized POLG transcript abundance was calculated as 2^⁻ΔCt^, where ΔCt = Ct(POLG) − Ct(Actb).

### Mitochondrial DNA quantification

Relative mitochondrial DNA (mtDNA) abundance was measured by duplex TaqMan qPCR on total DNA using TaqMan assays targeting the mitochondrial 12S rRNA gene (mt-Rnr1, Mm04260177_s1, Thermo Fisher) and the nuclear β-actin gene (*Actb*, Mm00607939_s1, Thermo Fisher). For each mouse, duplicate Ct values were averaged for each target, and ΔCt was calculated as Ct(*mt-Rnr1*) − Ct(*Actb*). Relative mtDNA abundance was calculated as 2^−ΔCt^ and expressed as a percentage of the mean 2^−ΔCt^ values in the control group.

### SDS-PAGE and immunoblotting

For SDS-PAGE, proteins were extracted from pulverized tissues using RIPA buffer (Thermo Fisher Scientific) supplemented with Complete protease inhibitor (Roche). After incubation on ice for 30 min, the samples were sonicated for 15 s using a tip sonicator. Following centrifugation at 16,000 x g for 20 minutes in a pre-chilled centrifuge, supernatants were collected, and protein concentration was determined using Bradford reagent (Thermo Fisher Scientific). 50-150 μg of protein extracts in Laemmli buffer were fractionated through Criterion polyacrylamide gels (Bio-Rad) and electroblotted onto PVDF membranes (Bio-Rad). Immunological detection was carried out using commercial antibodies: anti-POLG (ab128862, Abcam), anti-vinculin (ab129002, Abcam), anti-SDHA (ab14715, Abcam), anti-ATP5A (ab14748, Abcam), anti-UQCRC2 (ab14745, Abcam), anti-COXI (MT-CO1, ab14705, Abcam), anti-COXII (MT-CO2, ab110258, Abcam), anti-COXIV (11242-1-AP, Proteintech), anti-NDUFB8 (ab110242, Abcam) and anti-NDUFA13 (ab110240, Abcam).

### Immunofluorescence in tissues

Sagittal brain or transverse skeletal muscle sections (14 µm thick) were cut using a cryostat at −20°C (CM3050 S Leica Biosystems). The sections were either used immediately or stored at −80°C until analysis. Sections from mice of the same sex representing the different experimental groups were mounted on the same slide. For immunostaining, both brain and skeletal muscle sections were air-dried for 15 min. Skeletal muscle sections were further fixed with 4% paraformaldehyde (PFA) for 20 min in the dark. Next, the sections were permeabilized with 0.3% Triton X-100 in PBS for 10 min, followed by blocking with either 0.5% bovine serum albumin (BSA, Merck) and 2% goat serum (Merck) or a Mouse on Mouse IgG blocking reagent (Vector Laboratories) for 1 h. After a wash with PBS, the sections were incubated with different primary antibodies diluted in PBS containing 2% goat serum and 0.5% BSA overnight at 4°C. Following washing with PBS, the sections were incubated with Alexa Fluor-conjugated secondary antibodies (Thermo Fisher Scientific) for 1 h at room temperature. The following commercial antibodies were used: anti-calbindin (CB300, Swant AG), anti-Iba1 (019-19741, Wako Chemicals), anti-mCherry (AB0040-200, Cliniscience), anti-TFAM (19998-1-AP, Proteintech), anti-DNA (clone AC 30-10, Progen), anti-COXI (MT-CO1, ab14705, Abcam). For skeletal muscle, wheat germ agglutinin (WGA) conjugated to Alexa Fluor 488 (W11261, Thermo Fisher Scientific) was used to stain plasma membranes. Coverslips were mounted using DAPI-containing anti-fade mounting medium (Thermo Fisher Scientific) for nuclear staining. Images were acquired using a Leica SP5 confocal microscope with a 40× or 63× objective. To improve rendering, the brightness and contrast of the acquisitions were adjusted in Fiji ^25^.

### Cytochrome c oxidase (COX) and succinate dehydrogenase (SDH) activity staining

COX and SDH activity staining in brain and muscle sections was performed essentially as described previously ^20^. Sections from mice of the same sex representing the different experimental groups were mounted on the same slide. Briefly, for COX staining, frozen sagittal (brain) or transverse (muscle) sections were air-dried at room temperature for 15 minutes prior to incubation in freshly prepared COX staining solution (20 mg/mL catalase (Merck), 2 mg/mL reduced cytochrome c (Merck), and 1 mg/mL 3,3′-diaminobenzidine (DAB, Merck) in 50 mM sodium phosphate buffer, pH 7.4). Sections were rinsed in PBS and dehydrated through graded alcohols and xylene before mounting. For SDH staining, sections were incubated in a freshly prepared staining solution containing 1.5 mg/mL nitroblue tetrazolium (NBT, Merck), 130 mM sodium succinate (Merck), and 0.2 mg/mL phenazine methosulfate (PMS, Merck) in 0.05 M phosphate buffer, pH 7.6 (Merck) and processed as above prior to mounting. Stained sections were scanned at least 24 hours later using a NanoZoomer 2.0-HT slide scanner (Hamamatsu). Scans were visualized and processed using Hamamatsu’s NDP.view2 image viewing software.

### Hematoxylin and eosin (H&E) staining and estimation of Purkinje cell linear density

H&E staining was performed on sections of paraffin-embedded brain and liver after drying, deparaffinization, and rehydration. Each glass slide contained three sections from a single mouse brain. Nuclear staining was performed by incubation in hematoxylin for 10 min, followed by rinses in tap and distilled water. Cytoplasmic counterstaining was performed with alcoholic eosin Y for 1 min. Next, sections were briefly incubated in 70% ethanol (∼5 s), dehydrated in 80% and 100% ethanol (1 min each), and lastly in xylene. Slides were mounted using Pertex mounting medium and imaged with a NanoZoomer 2.0-HT (Hamamatsu Photonics). Scans were visualized and processed using Hamamatsu’s NDP.view2 image viewing software.

PC linear density was estimated in QuPath v. 0.7.0 ^26^ by counting all cells exhibiting characteristic dense H&E staining and/or a large soma within the entire length of the PC layer in two or three sagittal brain sections per mouse. To calculate PC density for each section, PC counts were divided by the PC layer length (mm) in that section, determined by drawing a polyline along it. Next, averaged PC densities per mouse were used to estimate the average PC densities of L/L and KO mice at a given age.

### Cell culture, transfection, and immunofluorescence

HeLa cells were seeded onto 13-mm coverslips in 6-well plates and transfected with 1 μg of TGPOLG1–9 plasmids using jetPRIME reagent (Polyplus-transfection). The following day, the cells were washed with PBS and fixed with 4% PFA in PBS for 10 min in the dark. Next, the cells were permeabilized for 10 min with 0.1% Triton X-100 (Merck) and blocked for 1 h at room temperature with blocking buffer containing 5% bovine serum albumin (Merck) and 0.1% Tween-20 (Merck) in PBS. The cells were then incubated for 1 h with anti-FLAG-M2 (F1804, Merck) and anti-Tom20 (11802-1-AP, Proteintech) antibodies, diluted in blocking buffer. Next, the coverslips were washed with PBS, and Alexa Fluor-conjugated secondary antibodies (Thermo Fisher Scientific) were added for 1 h at room temperature in the dark. Finally, the coverslips were mounted with anti-fade mounting medium (Thermo Fisher Scientific) containing DAPI for nuclear staining. Images were acquired using a Leica SP5 confocal microscope equipped with a 63×/1.40 NA oil-immersion objective. Brightness and contrast were adjusted in Fiji ^25^.

## Statistical analysis

Data were plotted and statistically analyzed in GraphPad Prism (version 11.1.0 (224)). The Holm–Bonferroni correction for multiple comparisons was performed manually. Statistical methods are specified in the figure legends, and only statistically significant differences are indicated in the figures.

## Results

### Loss of *Polg* in PV+ neurons causes a progressively worsening motor coordination deficit and a decreased lifespan

To demonstrate proof of concept for *POLG* gene replacement therapy, we first developed a conditionally targeted mouse model (*Polg^Lox^*^/Lox^) that allowed us to inactivate the *Polg* gene in PV+ neurons. Parvalbumin (PV) expression in rodents begins in the postnatal period and shows a broad temporal range across tissues ^27^. In the rodent CNS, parvalbumin exhibits a region-specific expression pattern: in the cerebellum, for example, PV expression is barely detectable during the first postnatal week but increases during the second to third postnatal weeks ^28^. The *PV-Cre* mouse line is often referred to as a neuronal recombination driver. However, parvalbumin is also expressed in skeletal muscle. Thus, *PV-Cre*-mediated recombination and *Polg* inactivation are expected not only in PV+ neurons but also in PV+ muscle fibers. To confirm and map *PV-Cre*-mediated recombination in the brain and skeletal muscle, we used a mouse line carrying a *Cre*-inducible tdTomato reporter whose expression, under normal conditions, is suppressed by a *lox*-flanked stop cassette upstream of the start codon ^22^. *Cre*-mediated recombination at the *lox* sites removes the stop cassette, enabling reporter expression. Immunofluorescence analyses of sagittal brain sections from 6-week-old *PV-Cre tdTomato* mice revealed strong tdTomato staining in the cerebellum, notably in Purkinje cells (PCs), as well as in individual neurons in the cortex and hippocampal regions (Figure 1A). Immunofluorescence analysis of tdTomato expression in three different muscle types (gastrocnemius, quadriceps, and soleus) showed that, as expected, different muscles contained varying proportions of tdTomato-positive muscle fibers, reflecting their relative fiber-type compositions: while quadriceps and gastrocnemius muscles contained many tdTomato-positive fibers, we detected no tdTomato expression in soleus muscle (Figure 1B). These data align with previous reports on the pattern of parvalbumin expression in skeletal muscle and suggest that *PV-Cre*-mediated inactivation of the *Polg* gene can also be expected in quadriceps and gastrocnemius muscles in *Polg* KO mice.

**Figure 1.**
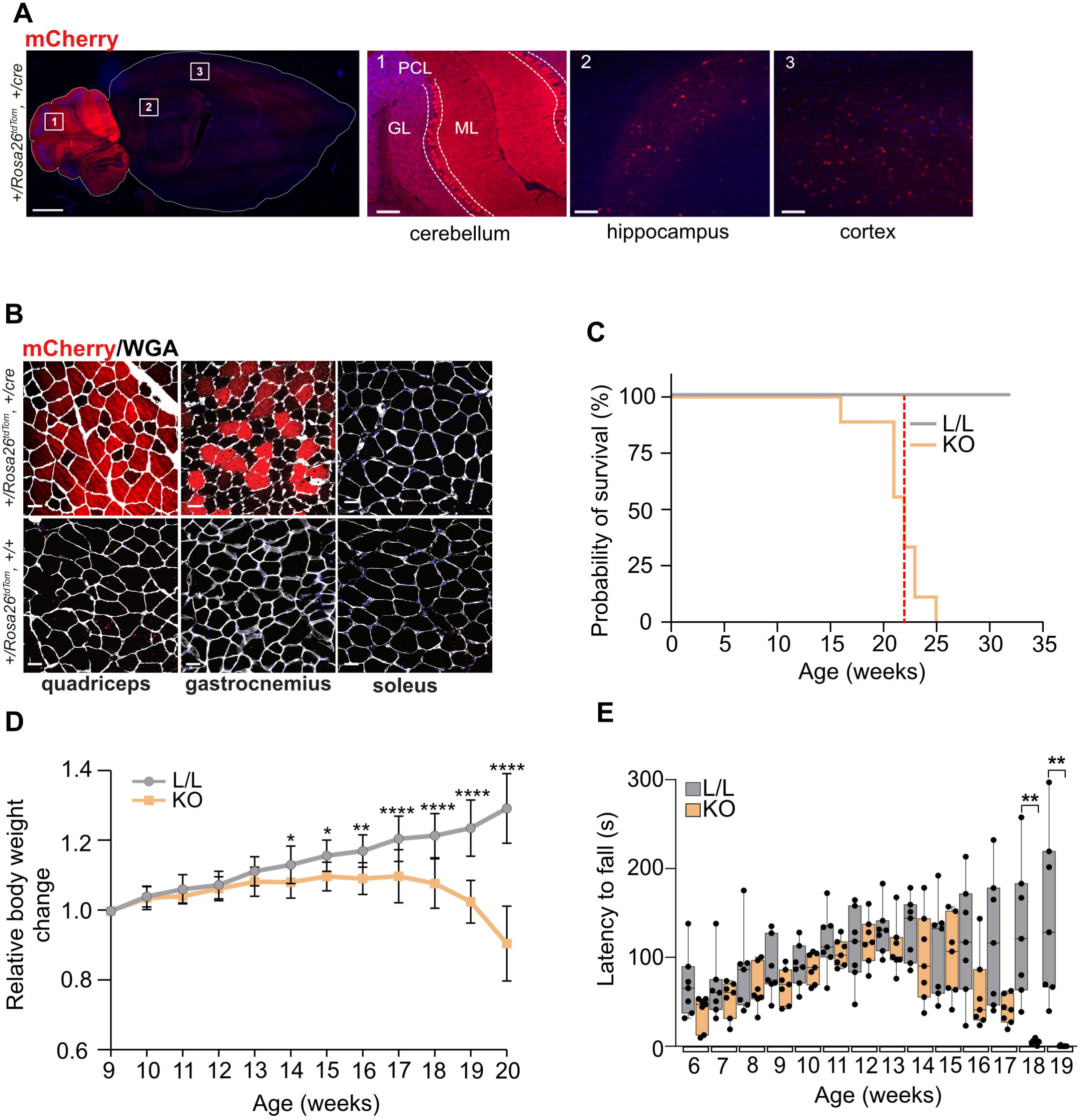
Mapping of *PV-Cre*-mediated recombination and characterization of *Polg*-deficient mice. **A.** Immunofluorescence analysis of tdTomato expression detected using an anti-mCherry antibody in brains from *PV-Cre*-expressing mice (*Rosa26^+/tdTomato^, +/PV-Cre*). Magnified images show PV+ neurons in the cerebellum, hippocampus, and cortical regions. Scale bars: whole-brain, 1 mm; magnified regions, 100 μm. PCL, Purkinje cell layer, delineated by dotted lines; ML, molecular layer; GL, granular layer. **B.** Immunofluorescence analysis of tdTomato expression in quadriceps, gastrocnemius, and soleus muscles in *PV-Cre*-expressing mice. *Rosa26^+/tdTomato^, +/+* mice did not express *PV-Cre* and were used as a negative control. Wheat germ agglutinin (WGA) staining delineates individual myofibers. Scale bar: 40 µm. **C.** Kaplan-Meier survival analysis of L/L and KO mice. Median survival of KO mice was 22 weeks (red dotted line). L/L, n = 6; KO, n = 9. **D.** Relative body weight change of L/L and KO mice at different ages. Body weight at each age was normalized to the body weight at 9 weeks of age. Data are presented as mean ± SD. L/L, n = 13; KO, n = 16. Mixed-effects analysis with Bonferroni correction, *, p < 0.05; **, p < 0.01; ****, p < 0.0001**. E.** Rotarod performance of L/L and KO mice at the indicated ages. Data are shown as box-and-whisker plots; boxes show the interquartile range with the median (line), and whiskers show the minimum and maximum values. Multiple Mann-Whitney tests with Bonferroni correction. L/L, n = 7; KO, n = 7. **, p < 0.01.

Consistent with the predominantly postnatal expression of *PV-Cre*, KO mice initially developed normally but later exhibited a progressively worsening disease. Median survival of KO mice was estimated at 22 weeks (Figure 1C). KO mice initially gained weight similarly to L/L littermates, but at ∼13-14 weeks of age, weight gain slowed, and they began to lose weight after ∼17 weeks of age (Figure 1D). At ∼19 weeks, KO mice developed tremors, had visibly impaired gait and overt motor coordination deficits. Therefore, given the disease trajectory and to prevent suffering, we established the humane endpoint for untreated mice at 19-20 weeks of age (terminal mice).

Next, we assessed the onset and progression of motor coordination deficits in KO mice using rotarod. This analysis measures the effects of neurological dysfunction on coordination, balance, and motor learning by recording how long a mouse remains on a rotating rod. Our analyses revealed that the latency to fall (LTF) of KO mice, while initially comparable to L/L littermates, decreased starting ∼16 weeks of age, and at 19 weeks, LTF was zero in all three repetitions for most tested KO mice (Figure 1E).

### Loss of *Polg* leads to RC deficiency in Purkinje cells

Loss of *Polg* led to a prominent motor coordination deficit, consistent with the observed reporter CNS expression pattern and suggesting cerebellar dysfunction. PCs are the sole output neurons of the cerebellar cortex, which is why we focused on these cells. H&E staining of brain sections from L/L and KO mice at 15 and 19-20 weeks of age revealed no major differences in the cerebellar layers or foliation, but a significant PC loss in terminal mice (Figure 2A). We also observed increased recruitment of Iba1-expressing microglial cells near PCs, suggesting a subtle neuroinflammatory response in terminal mice (Figure 2B).

**Figure 2.**
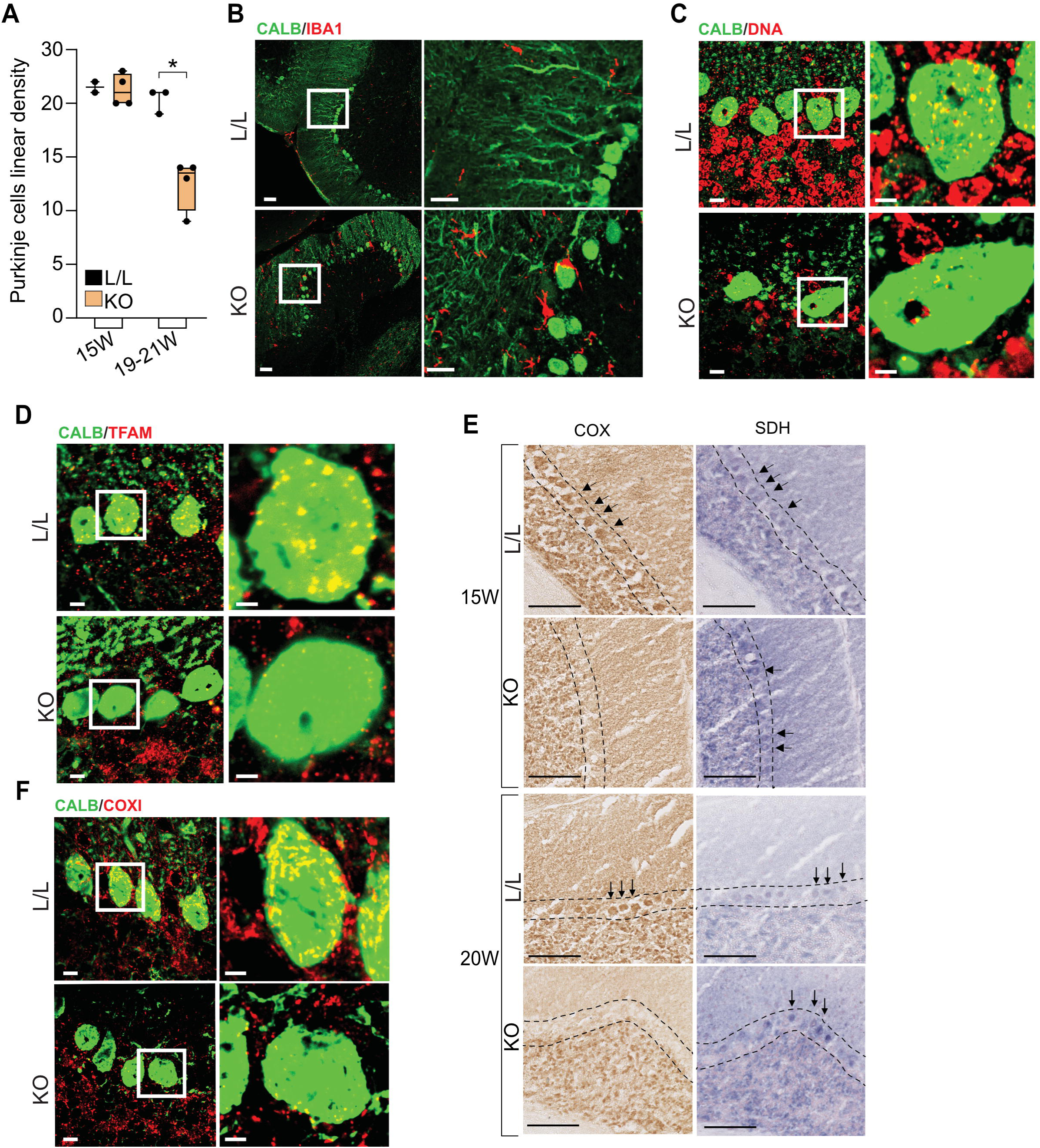
Purkinje cell loss and RC deficiency in *Polg* KO mice. **A.** Purkinje cell linear density (cells/mm) in L/L and KO mice at different ages. Data are presented as box-and-whisker plots; boxes indicate the interquartile range with the median (line) and whiskers indicate the minimum and maximum values. W, weeks of age; 15W L/L, n = 2; 15W KO, n = 4; 19-21W L/L, n = 3; 19-20W KO, n = 4. Welch’s t-test *, p < 0.05. **B.** Immunofluorescence analysis of cerebellar sections from 19-week-old L/L and KO mice. Purkinje cells were identified by calbindin immunostaining (green) and microglia by Iba1 immunostaining (red). Scale bars: 40 μm; magnified image, 20 μm. **C.** Immunofluorescence analysis of mtDNA foci in cerebellar sections from 19-week-old L/L and KO mice. Purkinje cells were immunostained as in B. DNA was detected using an anti-DNA antibody (red). Scale bars: 10 μm; magnified image, 5 μm. **D.** Immunofluorescence analysis of TFAM in cerebellar sections from 19-week-old L/L and KO mice. Purkinje cells were immunostained as in B. TFAM was detected using an anti-TFAM antibody (red). Scale bars as in C. **E.** COX and SDH activity staining in cerebellar sections from 15- and 20-week-old L/L and KO mice. The Purkinje cell layer is delineated, and arrows indicate individual PCs. Scale bar: 100 μm. **F.** Immunofluorescence analysis of COXI in cerebellar sections from 19-week-old L/L and KO mice. Purkinje cells were detected as in B (green). COXI was detected using a specific antibody (red). Scale bars are as in C.

To investigate the molecular consequences of *Polg* inactivation in PCs from KO mice, we assessed whether TFAM and mtDNA levels were affected using immunofluorescence. Because POLγA is critical for mtDNA maintenance, its loss causes mtDNA depletion and reduced abundance of the mtDNA packaging molecule and transcription factor TFAM ^29^. Along with other factors, mtDNA and TFAM form mitochondrial nucleoids that can be detected by immunofluorescence as DNA- or TFAM-positive foci throughout the PC soma ^30^. In PCs from KO mice, immunostaining with DNA- and TFAM-specific antisera revealed a marked decrease in both mtDNA- and TFAM-positive foci compared with L/L controls (Figure 2C, D). This suggests that *Polg* inactivation in PCs of KO mice resulted in mtDNA depletion.

Next, to assess the biochemical consequences of *Polg* inactivation, we performed enzymatic activity staining for cytochrome c oxidase (COX, respiratory chain complex IV) and succinate dehydrogenase (SDH, respiratory chain complex II) on brain sections from 15-week-old and terminal KO mice. This assay is commonly utilized in mitochondrial disease diagnostics to detect complex IV deficiencies arising from, for example, mtDNA gene expression defects ^31^. COX activity staining of sagittal brain sections from L/L mice revealed nearly uniform staining throughout most of the brain, making it difficult to distinguish individual cells. However, in the cerebellum, the large PC bodies stood out because of their strong COX activity staining (Figure 2E). In contrast, in the cerebellum of KO mice, PCs showed markedly reduced or even absent COX staining as early as 15 weeks of age, shortly before the onset of impaired rotarod performance. Conversely, SDH activity in individual PCs was detectable in KO mice at both ages. Together with the PC density measurements, this suggests that approximately half of PCs survive for at least several weeks after the onset of RC dysfunction. Moreover, immunofluorescent analysis revealed reduced COXI (an mtDNA-encoded subunit of complex IV) staining in KO PCs, corroborating the observed COX deficiency in these cells (Figure 2F). These results suggest a profound RC deficiency in cerebellar PCs, which likely underlies the motor coordination deficits of *Polg* KO mice.

Thus, we established a *Polg*-deficient mouse model that exhibits progressive motor coordination deficits and a shortened lifespan. In the CNS, this phenotype is associated with mtDNA depletion and RC deficiency in PCs. These findings allowed us to define therapeutic outcomes for our proof-of-concept strategy: (1) extended survival beyond 22 weeks and (2) rotarod performance at and beyond 19 weeks of age. At the tissue level, therapeutic outcomes included preservation of COX activity and nucleoid abundance in PCs. Moreover, we defined humane endpoints for treated mice primarily as the onset of overt gait or coordination abnormalities, with or without tremors and body weight loss exceeding 10%. Impaired rotarod performance alone was not an experimental endpoint.

### Development and characterization of gene therapy vectors expressing human POLG

In parallel, we designed nine gene therapy plasmids and tested their expression *in vitro* and *in vivo*. The large size of human *POLG* cDNA necessitated bespoke rAAV plasmids incorporating reduced-size regulatory elements ^20^. We utilized three ubiquitous promoters (miniCMV, EFS, and CAG) to control the expression of three *POLG* cDNA variants (wild-type, codon-optimized, and CpG-depleted), each fused to a C-terminal FLAG tag (Figure 3A). We confirmed expression and correct mitochondrial targeting of the POLγA-FLAG proteins *in vitro* by transfecting HeLa cells. Immunofluorescence analysis showed that all nine plasmids yielded POLγA protein that was correctly targeted to mitochondria (Figure 3B).

**Figure 3.**
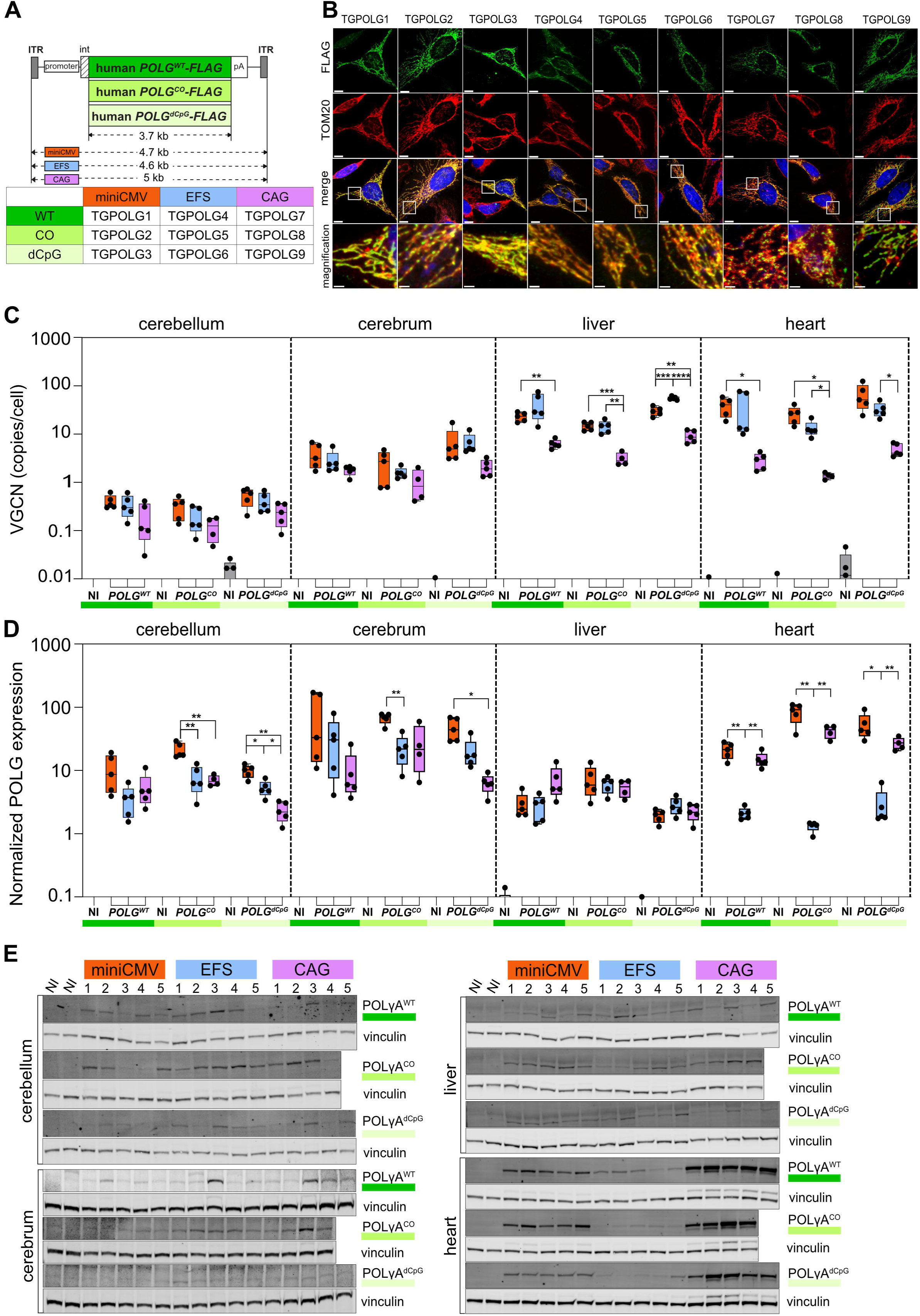
*In vitro* and *in vivo* characterization of nine *POLG*-expressing constructs. **A.** Top: schematic representation of the nine *POLG*-expressing constructs. The three *POLG* coding sequences are shown in different shades of green: *POLG^WT^* (dark green), *POLG^CO^* (light green), and *POLG^dCpG^* (pale green). The *POLG* cDNA and the vector genome lengths are indicated for each promoter-cDNA combination. Bottom: summary of the nine promoter-cDNA combinations. ITR, inverted terminal repeat; int, synthetic intron; pA, SV40 polyadenylation signal. **B.** Immunofluorescence analysis of HeLa cells transfected with TGPOLG1–9 plasmids. POLγA was detected using a FLAG-specific antibody (green). TOM20 was used as a mitochondrial marker (red). Scale bar: whole-cell image, 10 µm; magnified area, 2 µm. **C.** Vector genome copy number per cell in the indicated tissues of mice injected with TGPOLG1–9 vectors. The y-axis is displayed on a log10 scale. Data are presented as box-and-whisker plots; boxes indicate the interquartile range with the median (line) and whiskers indicate the minimum and maximum values. NI, non-injected, n = 2–5; TGPOLG1–9, n = 4–5. Welch’s ANOVA with Dunnett’s T3 multiple-comparisons test. Statistical comparisons were performed within each insert group. *, p < 0.05; **, p < 0.01; ***, p < 0.001; ****, p < 0.0001**. D.** RT-qPCR analysis of human POLG mRNA expression in the indicated tissues of mice injected with TGPOLG1–9. POLG expression was normalized to *Actb*. Data are presented as in **C**. NI, non-injected, n = 4–5; TGPOLG1–9, n = 4–5. Welch’s ANOVA with Dunnett’s T3 multiple-comparisons test. Statistical comparisons were performed within each insert group. *, p < 0.05; **, p < 0.01. **E.** Immunoblot analysis of different tissues from the mice above. Human POLγA protein was detected using an antibody specific to the human protein. Vinculin was used as a loading control. n = 4–5 mice per vector.

Next, we evaluated vector biodistribution across different tissues and their ability to drive *POLG* expression *in vivo*. To this end, we injected wild-type neonates (P0–P1) via the temporal vein with a single dose of each AAV9-pseudotyped vector at 1 × 10^14^ vg/kg. Systemic delivery via the temporal vein in neonates efficiently transduces both the CNS and different peripheral organs and tissues ^23,32^. We chose a high viral dose to compensate for vector dilution during growth, which may reduce transgene expression ^33,34^. At 9 weeks of age, we sacrificed all injected mice and collected their brains (separated into cerebrum and cerebellum), livers and hearts for downstream analyses.

First, we assessed vector biodistribution by measuring vector genome copy numbers (VGCN) of the different vectors across tissues (Figure 3C). These analyses showed that despite our early intervention, the CNS had the lowest transduction levels, with VGCN in the cerebellum being lower than in the cerebrum for the same vectors. The liver and heart showed comparable VGCN, at levels several-fold higher than in the CNS. The different vectors followed similar distribution patterns in the CNS, although the larger *CAG*-containing vectors tended to yield lower VGCN. This difference was even more evident in the liver and heart.

Next, we assessed the *POLG* mRNA expression levels in the different tissues by RT-qPCR (Figure 3D). These analyses showed that the miniCMV promoter generally drove higher expression levels in the CNS and heart, whereas all three promoters performed comparably in the liver. Interestingly, despite their low VGCN, CAG-containing vectors expressed more *POLG* in the heart, which may result from their well-known high activity in this tissue. Expression from EFS-containing vectors was intermediate in most tissues but lower in the heart compared to the other two vector groups.

Finally, to examine POLγA protein levels, we performed immunoblot analyses of whole-tissue extracts. Because the FLAG antibody could not detect the protein in whole-tissue lysates, likely due to low sensitivity, we used antisera that recognized human but not mouse POLγA (Figure 3E). This analysis showed that all nine vectors expressed POLγA protein in all examined tissues, with considerable variation in the heart. In this organ, despite lower VGCN, CAG-containing vectors drove higher mRNA and protein expression levels compared to the other rAAV variants. However, POLγA signals varied considerably among mice within each vector group, and some signals were near the detection limit of the antibody-based assay. As a result, we could not reliably quantify POLγA protein levels in these experiments.

These experiments demonstrate that AAV9-pseudotyped viral particles generated from all nine plasmids and injected into the temporal vein of neonates can transduce different tissues and drive immunodetectable CNS expression of POLγA *in vivo*.

### Single-dose temporal vein neonatal injection with *POLG-*expressing vectors delayed disease onset and extended the lifespan of KO mice

We established that all nine rAAV9-POLG vectors could drive expression of the human POLγA protein in the CNS. Next, we tested whether a single-dose injection in neonates could delay or prevent motor coordination deficits and extend the lifespan of KO mice. We focused on two lead candidates, TGPOLG4 and TGPOLG5. These vectors express the wild-type or codon-optimized *POLG* under the *EFS* promoter. We further modified TGPOLG4 and TGPOLG5 by removing the *FLAG* tag from the C-terminus of the *POLG* cDNA, yielding TGPOLG4.1 and TGPOLG5.1 (Figure 4A).

**Figure 4.**
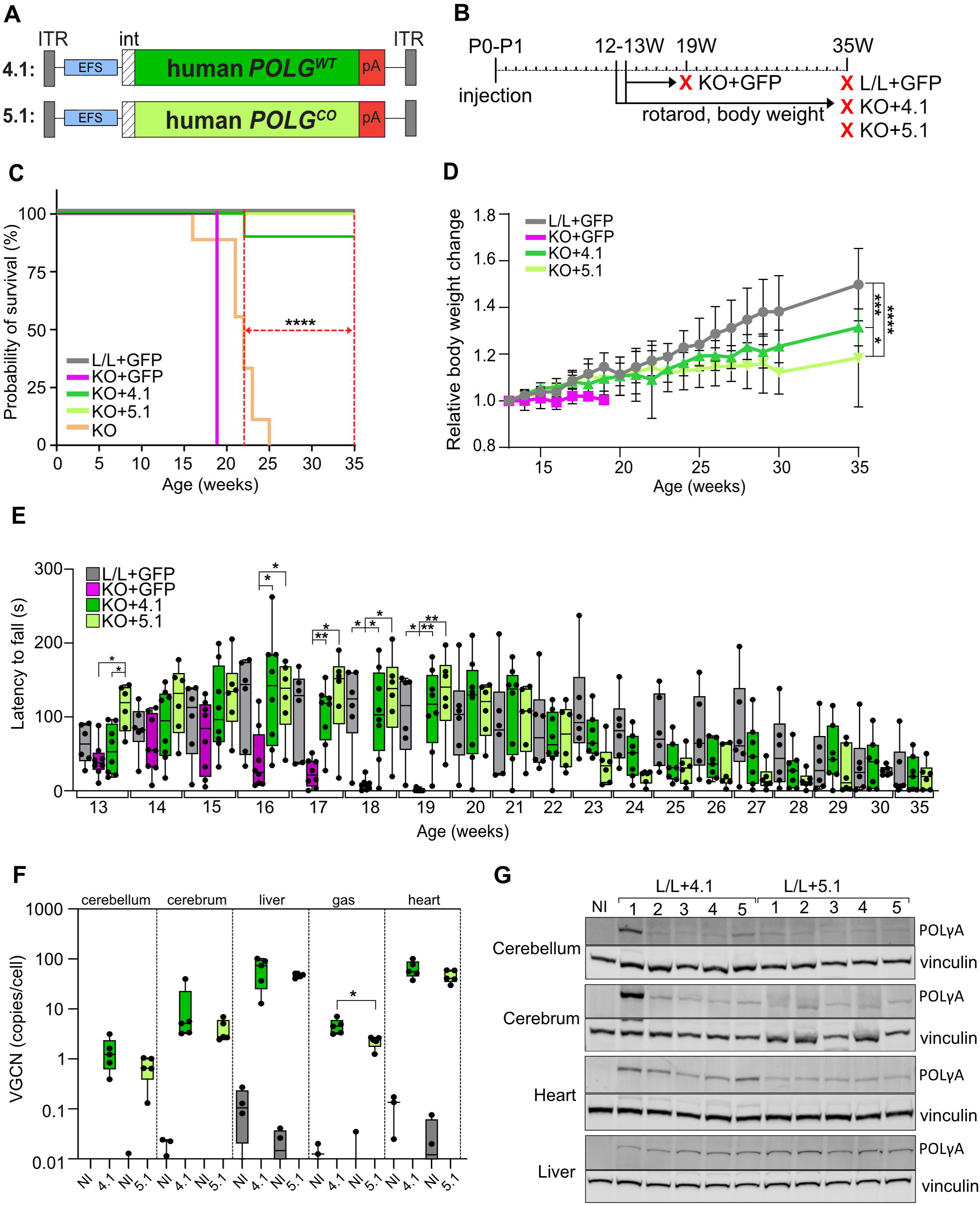
Neonatal administration of TGPOLG4.1 and TGPOLG5.1 delays the onset of motor coordination deficits and extends survival of *Polg* KO mice. **A.** Schematic representation of TGPOLG4.1 (4.1) and TGPOLG5.1 (5.1). ITR, inverted terminal repeat; int, synthetic intron; pA, polyadenylation signal. **B.** Experimental timeline. Vectors were administered at P0–P1. Analyses started at 12–13 weeks and continued until the end of the experiment at 35 weeks. KO + GFP mice were sacrificed at 19 weeks; surviving mice in the remaining groups were sacrificed at 35 weeks. **C.** Kaplan–Meier survival curves of the experimental groups. Untreated KO mice are shown for comparison. Red dotted lines indicate the median survival of untreated KO mice (left) and the end of the experiment (right). L/L + GFP, n = 6; KO + GFP, n = 8; KO + 4.1, n = 8; KO + 5.1, n = 6. Log-rank (Mantel–Cox) test; ****, *p* < 0.0001. **D.** Relative body weight change in the experimental groups. Data are shown as mean ± SD. Group sizes as in C. The KO + 4.1, KO + 5.1 and L/L + GFP groups at 35 weeks of age were compared. Mixed-effects analysis with Bonferroni’s multiple-comparisons test.*, p < 0.05; ***, *p* < 0.001; ****, *p* < 0.0001. **E.** Rotarod performance of the treatment groups at different ages. Data are presented as box-and-whisker plots: boxes indicate the interquartile range with the median (line) and whiskers indicate the minimum and maximum values. Group sizes are as in C. Two-way ANOVA with Tukey’s multiple-comparisons test. Comparisons are against KO + GFP. *, *p* < 0.05; **, *p* < 0.01. **F.** Vector genome copy numbers per cell in different tissues of TGPOLG4.1- and TGPOLG5.1-injected L/L mice. The y-axis is shown on a log10 scale. gas, gastrocnemius. Data are presented as in E. VGCNs of the two vectors is compared. Mann-Whitney tests. NI, non-injected, n = 3–4; 4.1, n = 5; 5.1, n = 5. *, *p* < 0.05. **G.** Immunoblot analysis of POLγA and vinculin in different tissues from TGPOLG4.1- and TGPOLG5.1-injected L/L mice. Group sizes as in F. NI, non-injected.

We injected neonates (P0–P1) with a single high dose (2×10^14^ vg/kg) of AAV9-pseudotyped TGPOLG4.1, TGPOLG5.1, or TG7-GFP vectors. Starting at 12-13 weeks of age, we monitored body weight and motor coordination as described above through the end of the experiment at 35 weeks of age (Figure 4B). Notably, 7/8 TGPOLG4.1- and 6/6 TGPOLG5.1-treated mice survived to 35 weeks of age, representing a significant lifespan extension compared to the median survival of KO mice (Figure 4C). As expected, GFP-injected KO mice exhibited the previously established disease trajectory: they failed to gain weight (Figure 4D) and developed progressively worsening motor coordination (Figure 4E). At 19 weeks of age, these mice had overt gait abnormalities, consistently failed the rotarod test (LTF ≤ 5 s, arbitrary threshold; 6/8 mice had LTF = 0) and were euthanized. In striking contrast, at this age, 7/8 TGPOLG4.1- and 6/6 TGPOLG5.1-injected KO mice had rotarod performance comparable to GFP-injected L/L controls and showed no overt signs of disease or distress (Figure 4E). Only one TGPOLG4.1-injected KO mouse developed a disease phenotype and was euthanized at 22 weeks of age. However, rotarod performance in TGPOLG4.1- and TGPOLG5.1-treated mice declined starting 23 weeks of age and remained lower than in the control group thereafter. In the control group, rotarod performance was initially higher but gradually declined as well. Weight gain, while initially comparable to controls, slowed after ∼22 weeks of age (Figure 4D). At 35 weeks of age, 1/6 mice in the control group, 3/7 mice in the TGPOLG4.1-injected group, and 2/6 mice in the TGPOLG5.1-injected group had LTF < 5 s. Despite low rotarod performance, 35-week-old treated KO mice showed no overt signs of disease (tremors, impaired gait or overt coordination deficits) and, aside from being leaner, were otherwise indistinguishable from controls.

We assessed vector biodistribution and POLγA protein levels in tissues from 35-week-old TGPOLG4.1- and TGPOLG5.1-injected L/L mice. At this age, VGCN followed the previously established pattern for the parent vectors, TGPOLG4 and TGPOLG5, with the lowest levels in the cerebellum, intermediate levels in the cerebrum, and highest levels in the liver and heart (Figure 4F). In this analysis, we also included gastrocnemius muscle, which showed intermediate VGCN comparable to those in the cerebrum. Immunoblot analyses of whole tissue extracts showed immunologically detectable levels of POLγA in all examined tissues except gastrocnemius (Figure 4G). Thus, injection with TGPOLG4.1 and TGPOLG5.1 during the neonatal period results in sustained vector persistence and POLγA expression in different disease-relevant tissues.

### Expression of human *POLG* preserved COX activity and mtDNA levels in Purkinje cells of KO mice

Loss of *Polg* decreased nucleoid foci abundance in untreated KO mice. Therefore, we assessed whether a single-dose injection with TGPOLG4.1 or TGPOLG5.1 preserved PC nucleoids in treated KO mice at 35 weeks of age. Immunofluorescence analysis with DNA- and TFAM-specific antisera showed that nucleoid foci were barely detectable in 19-week-old GFP-injected KO mice but were clearly preserved in 35-week-old KO mice treated with TGPOLG4.1 or TGPOLG5.1 (Figure 5A, B). Consistent with these data, COX activity staining and COXI protein levels, while decreased in PCs from GFP-injected KO mice, were preserved in KO mice treated with either therapeutic vector (Figure 5C, D).

**Figure 5.**
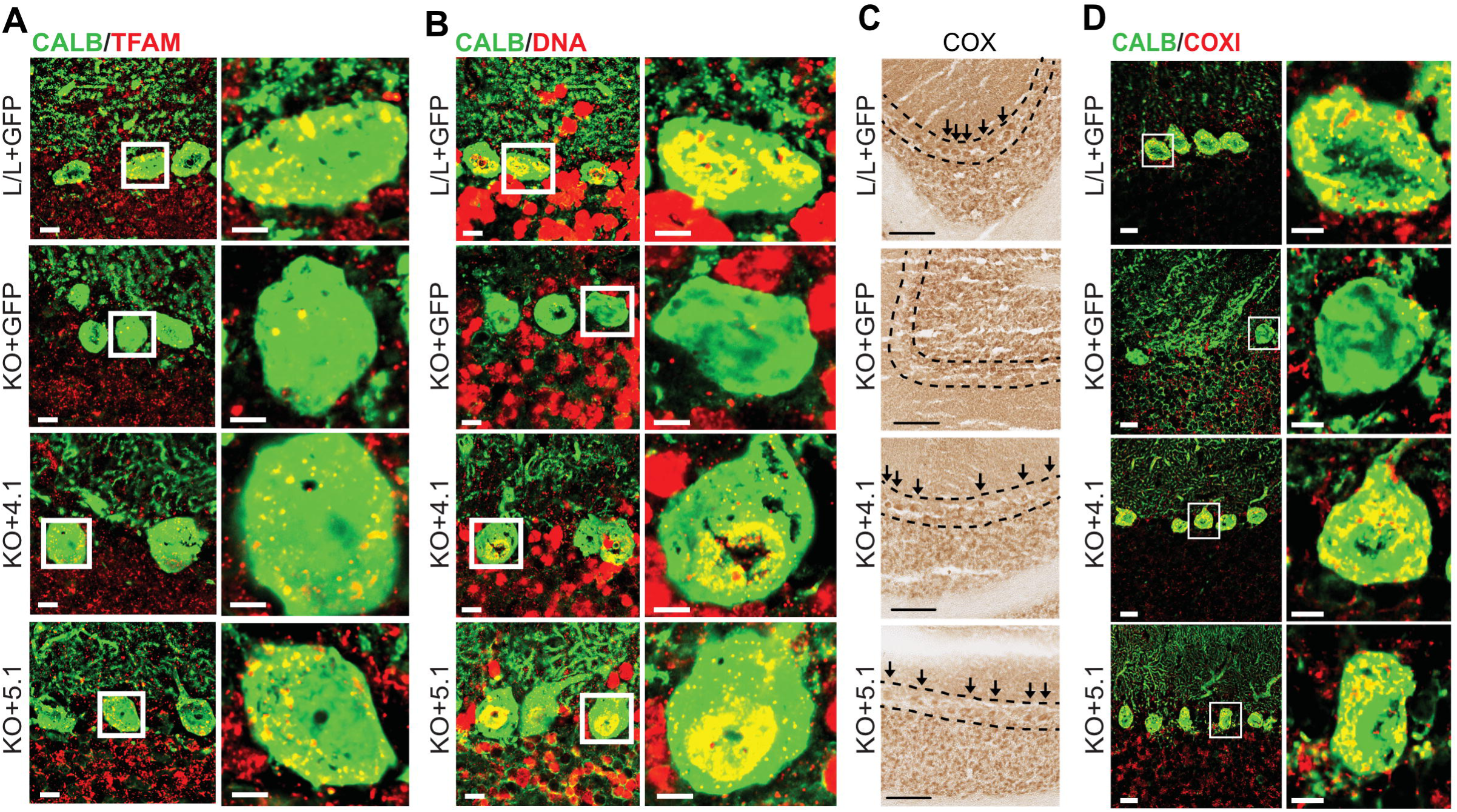
*POLG* expression preserves mtDNA foci abundance and COX activity in PCs of KO mice. **A.** Immunofluorescence analysis of brain sections from GFP-injected L/L mice and KO mice injected with either GFP, TGPOLG4.1 or TGPOLG5.1. Purkinje cells (green) and TFAM (red) were detected as in Figure 2D. Scale bars: 10 μm; magnified image, 5 μm. **B.** Immunofluorescence analysis of brain sections from mice in the different treatment groups (see A). Purkinje cells (green) and DNA (red) were detected as in Figure 2C. Scale bars as in A. **C.** COX histochemical staining of brain sections from mice injected with the different vectors (see A). Dotted lines delineate the Purkinje cell layer, and arrows indicate individual Purkinje cells. Scale bar: 100 μm. **D.** Immunofluorescence analysis as in A. Purkinje cells (green) and COXI (red) were detected as in Figure 2F. Scale bars as in A.

Thus, neonatal *POLG* gene replacement therapy preserved mtDNA foci abundance and COX activity in PCs, delayed the onset of motor coordination deficits, and extended survival of *Polg* KO mice.

### Expression of human *POLG* preserved COX activity, RC subunit abundance and mtDNA levels in the skeletal muscle of KO mice

Our tdTomato reporter mapping showed that *PV-Cre* is active in some quadriceps and gastrocnemius muscle fibers. This strongly suggests that *Polg* inactivation also occurs in these skeletal muscle types in KO mice. Therefore, we performed COX and SDH activity staining on transverse quadriceps muscle sections from L/L, untreated, and treated KO mice (Figure 6A). In L/L mice, COX staining intensities were notably heterogeneous across muscle fibers, likely reflecting differences in mitochondrial content. SDH staining supported this, showing similar heterogeneity throughout the muscle section. In muscle sections from untreated KO mice, we observed a pronounced decrease in COX activity staining in some muscle fibers, while the SDH staining pattern remained similar between KO and L/L mice. This finding suggests a complex IV deficiency in individual muscle fibers. In contrast, COX activity in quadriceps muscles from treated mice was variably preserved at 35 weeks of age. Immunoblot analysis of different RC subunits further corroborated this result (Figure 6B, C). Levels of complex IV subunits COXI, COXII, and COXIV, and complex I subunits NDUFB8 and NDUFA13, were decreased in the quadriceps muscle of KO mice. In contrast, levels of the nucleus-encoded SDHA (a subunit of complex II) and ATP5A (a subunit of complex V) were unchanged. Treatment with either therapeutic vector preserved complex I and IV subunit levels, albeit to varying degrees in different mice. The pattern of changes in subunit abundance is consistent with defects in multiple RC complexes and with mtDNA depletion following *Polg* inactivation. Indeed, relative mtDNA abundance was lower in KO mice than in controls, whereas it was variably preserved after treatment with either gene replacement vector (Figure 6D).

**Figure 6.**
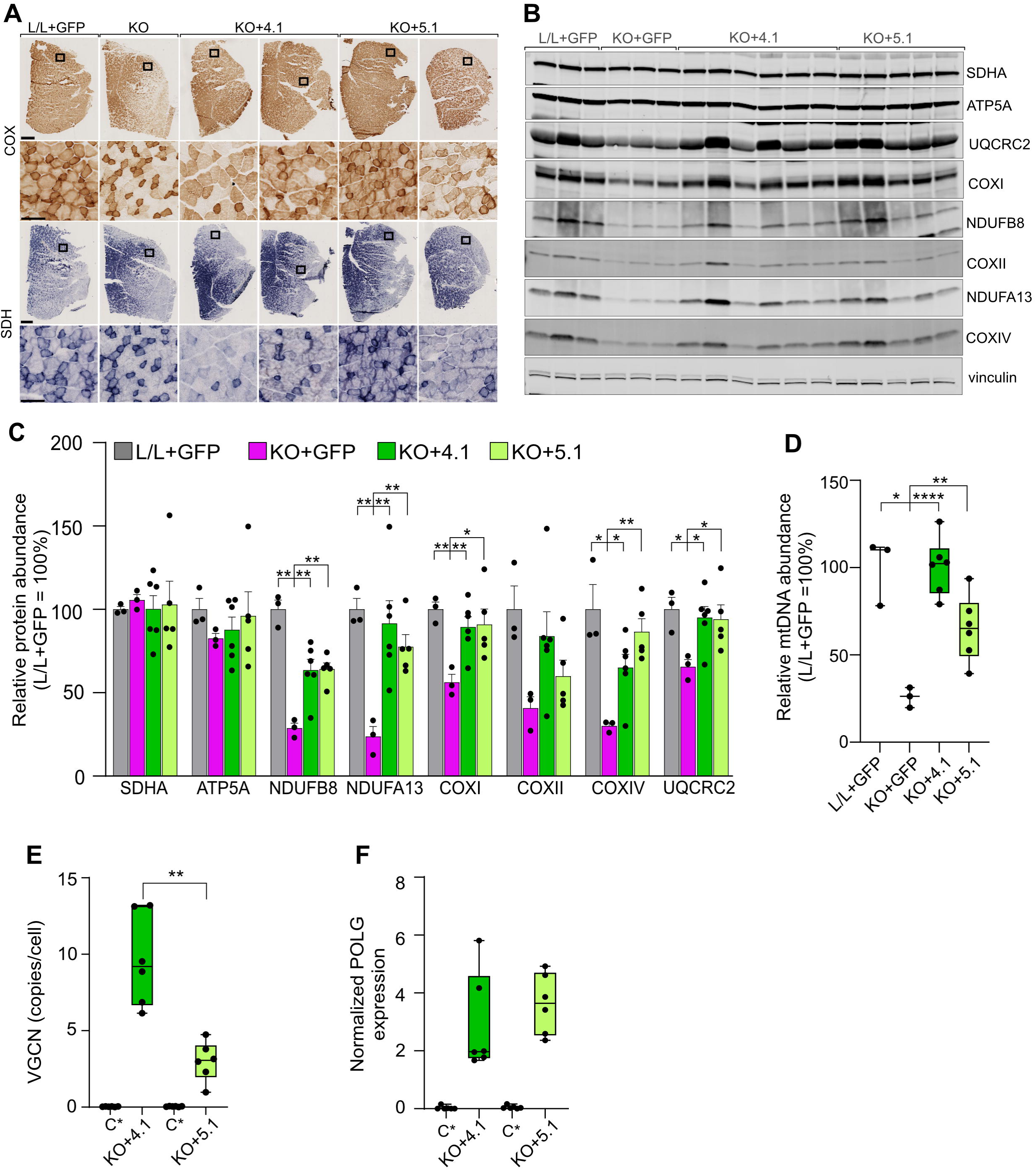
*POLG* expression preserves COX activity and steady-state levels of oxidative phosphorylation subunits in quadriceps muscle of KO mice. **A.** COX and SDH staining of quadriceps sections from 35-week-old mice in the indicated treatment groups. Untreated KO mice (19 weeks old) were used for comparison. Scale bars: whole-muscle section, 500 µm; magnified area, 100 µm. **B.** Immunoblot analysis of subunits from the five oxidative phosphorylation complexes and vinculin in quadriceps muscle from the different experimental groups. **C.** Quantification of (B). Band intensities were normalized to vinculin, averaged, and shown relative to L/L + GFP. Data are presented as mean ± SEM. Statistical comparisons were against KO + GFP in each protein group. L/L + GFP ,n = 3; KO + GFP, n = 3; KO + 4.1, n = 6; KO + 5.1, n = 5. Multiple Welch’s t-tests with Holm–Bonferroni correction for multiple comparisons. *, *p* < 0.05; **, *p* < 0.01. **D.** Relative mtDNA abundance in the indicated treatment groups. L/L + GFP, n = 3; KO + GFP, n = 3; KO + 4.1, n = 6; KO + 5.1, n = 6. Each group was compared against KO + GFP. Welch’s ANOVA with Dunnett’s T3 multiple-comparisons test. *, *p* < 0.05; **, *p* < 0.01; ****, *p* < 0.0001. **E.** VGCN per cell in quadriceps muscle from the indicated experimental groups. C*, pooled VGCN measurements for GFP-injected L/L (n = 3) and KO (n = 3) mice; 4.1, n = 6; 5.1, n = 6. Welch’s t-test. **, *p* < 0.01. **F.** RT-qPCR analysis of human POLG mRNA expression in quadriceps muscle from the different experimental groups. *POLG* expression was normalized to *Actb*. Group sizes were as in E. Data in **D, E** and **F** are presented as box-and-whisker plots where boxes indicate the interquartile range with the median (line) and whiskers indicate the minimum and maximum values.

VGCN analysis showed that TGPOLG4.1 levels were generally higher than TGPOLG5.1, with considerable variation between individual mice (Figure 6E). We were unable to detect the POLγA protein by immunoblotting. However, RT-qPCR analysis demonstrated that the two *POLG* cDNA variants were expressed in quadriceps muscle (Figure 6F). Combined with the clear preservation of RC subunits in treated mice, this strongly suggests the presence of functional POLγA protein.

Cumulatively, these results suggest that rAAV9-TGPOLG4.1 and rAAV9-TGPOLG5.1 can deliver and drive expression of the human POLγA protein in skeletal muscle, another POLG disease-relevant tissue.

## Discussion

Here, we demonstrate a preclinical proof of concept for *POLG* gene replacement therapy, in which neonatal injection with therapeutic vectors expressing human *POLG* delayed the onset of motor coordination deficits and extended the lifespan of *Polg*-deficient mice. This early-life intervention preserved RC complex IV activity in PCs, whose dysfunction partly underlies the clinical phenotype of untreated mice. In skeletal muscle of untreated *Polg* KO mice, we observed decreased abundance of subunits from complexes I and IV, suggesting a multiple RC deficiency. Administration of the therapeutic vectors preserved COX activity and the abundance of individual RC subunits. These results, together with persistent *POLG* expression in another disease-relevant tissue, the liver, demonstrate multi-tissue delivery and support the feasibility of developing a gene therapy for *POLG* disease.

We report two advances that enable further preclinical development of *POLG* gene replacement therapy. First, we developed a conditional *Polg* knockout in PV+ neurons and skeletal muscle to serve as a preclinical platform for gene therapy development. PV+ neurons include several distinct neuronal populations, most notably fast-spiking inhibitory interneurons in the cerebral cortex and hippocampus, PCs and other inhibitory neurons in the cerebellum, and proprioceptive neurons in the dorsal root ganglia ^28,35,36^. Collectively, they participate in the timing and activity of neuronal circuits and contribute to sensory processing, network synchronization, proprioception and motor coordination. In patients with Alpers syndrome, PV+ inhibitory neurons in the occipital cortex and cerebellar PCs are particularly sensitive to RC dysfunction, and their loss contributes to epilepsy and ataxia in these patients ^37,38^. However, epilepsy and ataxia (cerebellar and sensory) are not limited to Alpers syndrome and are recognized *POLG* disease presentations. Studies of *PV-Cre*-mediated *Tfam* and *Frataxin (Fxn)* KO mice have shown that mitochondrial dysfunction in PV+ neurons leads to strong neurological phenotypes, including sensory and/or cerebellar ataxia, seizures or seizure-like behavior, and reduced survival ^35,36^. RC dysfunction was detected in PV+ interneurons in the thalamic reticular nucleus, and dysfunction in this region was tentatively linked to the stargazing episodes. Our *Polg* KO mice developed a molecular and behavioral phenotype resembling that of the *Tfam* KO mice with mtDNA depletion and RC deficiency in PCs accompanied by a progressively worsening motor coordination deficit consistent with cerebellar dysfunction. Although we focused on PCs, loss of *Polg* could also affect PV+ neurons in the thalamus and visual cortex, but whether this leads to seizures needs to be demonstrated. Nevertheless, the progressive neurological phenotype and the underlying mtDNA depletion and RC defects establish the relevance of our model as a therapeutic platform for POLG disease. Importantly, our proof of concept shows that neonatal AAV9-*POLG* gene therapy can preserve mtDNA levels and RC function in cerebellar PCs and delay the development of severe neurological disease. This establishes the feasibility of *POLG* gene replacement therapy to treat cerebellar manifestations in *POLG* disease.

Using the tdTomato reporter, we demonstrated that *Cre* recombination also occurs in individual gastrocnemius and quadriceps muscle fibers, likely leading to *Polg* loss *in KO mice*. Consistent with this, we detected a multiple RC defect affecting complex I and complex IV in quadriceps muscle, strongly supporting muscle involvement in *KO* mice and further strengthening our model’s relevance to *POLG* disease. Skeletal muscle involvement is a well-recognized disease presentation in humans, affecting both pediatric and adult patients ^10^. It can present as isolated proximal or distal weakness and exercise intolerance, ptosis and progressive external ophthalmoplegia (PEO). Muscle biopsies in patients with *POLG*-linked myopathy frequently reveal COX-deficient and ragged-red fibers, multiple mtDNA deletions or depletion ^39^. Importantly, we demonstrated that neonatal AAV-*POLG* administration preserved COX activity and the abundance of CI and CIV subunits. This strongly supports the feasibility of our *POLG* gene replacement therapy approach for treating *POLG*-linked myopathy.

The second important advance was demonstrating a single-vector approach for *POLG* gene therapy. *POLG* cDNA was too large to include in classical rAAV, as it would exceed the optimal packaging capacity of the AAV capsid (∼4.7-5 kb). Instead, we used a smaller rAAV plasmid design that lacked part of the noncoding rAAV sequence and included minimal promoters and a shorter SV40 poly(A) signal ^20^. Thus, the miniCMV- and EFS-containing rAAV-POLG vectors remained at or below the upper viral packaging limit. The CAG-containing plasmids were slightly larger because they used the conventional-size CAG promoter, leading to reduced viral production titers (not shown) and an overall tendency for lower VGCN in all examined tissues. This aligns with observations that, even with minimally oversized vectors, the proportion of truncated genomes increases in the viral preparations. Nevertheless, all tested vectors targeted the CNS and expressed the *POLG* transgene. Expression levels varied, with the miniCMV promoter driving higher CNS expression than the other two promoters. In the liver, expression was largely comparable. Only in the heart did the EFS promoter show lower activity than the miniCMV and CAG promoters. In the liver and heart, CAG-containing vectors, despite lower VGCN, were associated with mRNA levels similar to or higher than those of the other two vector groups. However, in the CNS, VGCN and *POLG* mRNA levels associated with the CAG-containing vectors followed similar trends. While the miniCMV promoter outperformed the other two, it should be noted that its full-length equivalent has been shown to undergo tissue- and cell type-dependent silencing *in vivo*, including in the liver, spinal motor neurons, dorsal root ganglion neurons, and several CNS regions ^40,41^. Whether the miniCMV promoter is also silenced in these tissues is unclear. Such silencing would be undesirable for sustained long-term transgene expression and warrants further evaluation. In this proof of concept, we focused on two vectors carrying EFS promoters, which showed consistent expression levels in the CNS, liver and skeletal muscle (three *POLG* disease-relevant tissues), but were least active in the heart. The relevance of this finding for the feasibility of POLG gene replacement therapy with these vectors is unclear. Cardiomyopathy is extremely rare in *POLG* disease and is not present in our model, preventing us from determining whether POLG expression from these vectors was sufficient to maintain RC function in the heart. Additional studies will address this question and further characterize the short- and long-term therapeutic potential of the remaining rAAV-POLG vectors. For now, we have established a toolbox of rAAV-POLG vectors with ubiquitous promoters that can target the CNS and express human *POLG* cDNA in different tissues.

To ensure efficient brain targeting, we chose the AAV9 capsid and a high injection dose. The AAV9 capsid is well characterized and has broad tissue tropism, targeting many of the tissues affected in POLG disease ^42^. Moreover, AAV9 is already used in the FDA-approved Zolgensma and Itvisma gene replacement therapies for spinal muscular atrophy. However, because AAV9 is a naturally occurring viral capsid, some patients may already have pre-existing immunity that can neutralize the vector and prevent transgene expression ^43,44^. The likelihood of developing such immunity increases with age, rendering some juvenile/adult *POLG* patients ineligible for AAV-based POLG gene replacement therapy. Immune cross-reactions between capsids can further increase the risk of vector neutralization, even in patients with low pre-existing anti-AAV9 immunity. Interestingly, AAV9 seropositivity has also been reported in newborns due to maternally transferred antibodies, but in these cases, antibody titers usually decrease in the following months ^45^. Avoiding pre-existing immunity may be possible by using engineered capsids with altered antigenicity or potentially using nonviral delivery platforms ^46,47^. Alternatively, in neurological diseases, direct intracranial or intrathecal delivery may help reduce exposure to pre-existing immunity by partially shielding the vector from circulating antibodies ^48^.

High viral doses are associated with safety concerns. In clinical trials, these have been associated with serious adverse events, such as hepatotoxicity, severe immune responses, and, rarely, death ^44,49^. Here, we did not observe any increased mortality or overt disease in injected neonates. Lowering the viral dose helps reduce the risk of toxic side effects but may substantially decrease therapeutic efficacy. A dose de-escalation study will therefore help determine the minimum effective systemic dose required to achieve therapeutic benefit in *Polg* KO mice. The minimum effective dose is likely to be tissue-specific and to depend on the lowest POLγA level required to maintain mitochondrial function in that tissue. Fully restoring endogenous POLγA levels is probably not required. Heterozygous *Polg* KO mice express only ∼50% of the wild-type transcript, yet their mtDNA levels are comparable to or slightly lower than those in wild-type mice, and they develop normally ^50^. Moreover, heterozygous carriers of recessive null or splice variants in the *POLG* locus are asymptomatic ^51,52^. *POLG* overexpression, on the other hand, is well tolerated in cultured mammalian cells, although this has not been tested *in vivo*. Only in *Drosophila*, supraphysiological levels (∼50-fold higher than endogenous) of the *pol γ-α* subunit produced mtDNA depletion and developmental defects ^53^. However, *POLG* expression from the vectors developed here is unlikely to reach such high levels in most targeted tissues because minimal promoters appear less active than their full-length equivalents, although this can be tissue-specific or depend on the vector design ^54,55^. A possible exception could be the liver, where AAV accumulation may increase hepatocyte transduction, leading to POLG overexpression. In turn, this may exacerbate vector-induced hepatic immune or cytotoxic responses. We did not assess these effects, although *POLG* mRNA expression levels in the liver were comparable to those in the cerebrum and cerebellum.

Here, we expressed *POLG* in presymptomatic mice, demonstrating the disease-preventive benefit of our approach. However, it remains to be seen whether post-symptomatic administration can arrest disease progression. Interestingly, we observed COX-deficient PCs as early as 15 weeks of age, yet they remained identifiable and SDH-positive for at least another 5 weeks, well after disease onset. This raises the possibility that re-expression of POLG in these cells may restore RC biogenesis and function. This is consistent with findings in *Tfam* KO and *Fxn* KO mice, showing that PV+ neuronal cells, including PCs, can endure mitochondrial dysfunction for some time before irreversible cell loss. Indeed, AAV-*FXN* administration in post-symptomatic *Fxn* KO mice rescued their motor coordination deficits ^36^. In a clinical setting, where POLG disease has already progressed before diagnosis, re-expression of functional *POLG* will not reverse cell loss, but it can prevent further damage and arrest disease progression. The extended interval between *Polg* inactivation and the onset of severe motor coordination deficits in our model provides a window of opportunity to test this hypothesis.

Previous studies have shown that defects in mtDNA maintenance are amenable to AAV-mediated gene replacement. Liver-directed AAV2/8-MPV17 restored mtDNA copy number and RC function, and prevented ketogenic diet-induced liver failure in *Mpv17*-deficient mice ^56^. Similarly, neonatal systemic delivery of AAV9-DGUOK in *Dguok-*deficient mice restored liver mtDNA levels, improved RC complex activities, ameliorated liver dysfunction, and prolonged survival ^57^. In *Tk2*-deficient mice, AAV-mediated gene replacement restored TK2 activity in most examined tissues, delayed disease onset and prolonged survival ^58^. Notably, AAV-TK2 therapy was potentiated by dC/dT supplementation, an FDA-approved treatment for TK2 disease ^59^. This illustrates the rationale for combined pharmacological and AAV treatments. Currently, dC/dT supplementation is in an open-label, single-arm Phase II trial for *POLG* disease ^60,61^, while a small-molecule POLγ activator, PX578 (Pretzel Therapeutics), is expected to enter a Phase II clinical trial. Unlike PX578, which acts directly on the POLγ enzyme, enhancing its activity and processivity, dC/dT supplementation aims to stimulate POLγ activity and mtDNA replication by increasing substrate availability ^62,63^. Beyond their therapeutic promise as standalone therapies, these treatments could also play a supportive or potentiating role in *POLG* gene therapy. In one scenario, pharmacological treatment can extend the therapeutic window for gene therapy, especially in neonates with severe disease. In another, not mutually exclusive scenario, it can accompany gene replacement therapy, especially for multisystemic disease affecting difficult-to-transduce organs or when lower AAV-*POLG* doses are used, helping achieve better and longer-lasting therapeutic outcomes.

In summary, our study provides the first proof of concept for AAV-mediated *POLG* gene replacement therapy. We developed a disease-relevant mouse model and a series of AAV vectors carrying POLG cDNA under the control of ubiquitous promoters. We showed that these vectors can deliver the *POLG* cDNA and drive its expression in different disease-relevant tissues, providing a demonstrable therapeutic benefit in two cases. Our work lays the foundation for further development of POLG gene replacement therapy and its clinical translation.

## Acknowledgements

M.D.M. and B.R. thank Julie de Luxembourg, Robert de Luxembourg, and Dr. Brian S. Tseng for their support throughout the project. M.D.M. thanks Dr. Agnès Rötig for providing access to laboratory space and equipment. We thank the Histology Core Facility (SFR Necker, Inserm US24–CNRS UAR 3633); ARC for providing access to the NanoZoomer 2.0 HT (Hamamatsu); Dr. Meriem Garfa Traore for help with cell imaging; and the staff of the animal facility (LEAT) at the Imagine Institute. We acknowledge the Imagine Institute’s rAAV platform for producing the viral preparations used in this study.

This work is dedicated to Frederik de Nassau and to all individuals affected by POLG disease and their families.

## Author contributions

**S.Z., S.M., E.A., M.T.P.-C. and C.G.:** Data acquisition and analysis, writing: review & editing. **M.S.S.:** AAV production, writing: review & editing. **S.C.K. and S.A.M.:** mouse model generation, preliminary characterization, writing: review & editing. **A.M.:** funding acquisition, writing: review & editing. **B.R.**: conceptualization, funding acquisition, writing – original draft, review & editing. **M.D.M.:** conceptualization, project supervision, data curation and analysis, funding acquisition, writing – original draft, review & editing.

## Data availability

All data are available within this article. Requests for resources and reagents should be addressed to Dr. Metodi Metodiev.

